# Iron deprivation activates aboveground cell wall biosynthesis in *Populus* and the role of PtrbHLH011

**DOI:** 10.1101/2024.06.28.601228

**Authors:** Dimiru Tadesse, Yuqiu Dai, Lin Yang, Yang Yang, Nidhi Dwivedi, Desigan Kumaran, Crysten E. Blaby-Haas, Anna Lipzen, Kassandra Santiago, Kerrie Barry, Chang-Jun Liu, Meng Xie

**Affiliations:** Biology Department, Brookhaven National Laboratory, Upton, NY, 11973, USA; National Synchrotron Light Source, Brookhaven National Laboratory, Upton, New York, NY, 11973, USA; Department of Energy Joint Genome Institute, Lawrence Berkeley National Laboratory, Berkeley, CA, 94720, USA; Molecular Foundry, Lawrence Berkeley National Laboratory, Berkeley, CA, 94720, USA

## Abstract

Lack of mechanistic understanding of the environmental plasticity of secondary cell wall (SCW) biosynthesis hinders the massive bioenergy production on marginal lands. Growing bioenergy crops on marginal lands is appealing to minimize competition for arable land. However, abiotic stresses, particularly iron deficiency stress, are widespread to perturb SCW biosynthesis. In poplar, a major bioenergy crop, we demonstrated that iron deprivation activates stem SCW biosynthesis and identified transcription factor PtrbHLH011 as a possible underlying regulator. PtrbHLH011 is a potent repressor of SCW, whose overexpression resulted in a reduction of stem SCW by over 65%. Our genomic and molecular studies discovered that PtrbHLH011 binds to the AAAGACA sequence and represses essential genes for SCW biosynthesis, flavonoid biosynthesis, and iron homeostasis. Wood formation and iron deprivation downregulates PtrbHLH011 to release the transcriptional repression. Our findings reveal a regulatory mechanism coordinating SCW biosynthesis in response to environmental iron availability and suggest that PtrbHLH011 manipulation may help engineer bioenergy crops with improved performance under marginal conditions.

## Introduction

*Populus* (poplar) is an important renewable resource for lignocellulosic biomass-based bioenergy production. In addition, poplar is a well-established model system for studying woody perennial plant-specific traits, such as wood formation, secondary growth, and adaptation to environmental change (Wullschleger et al., 2013). Lignocellulosic biomass is produced through secondary growth and secondary cell wall (SCW) biosynthesis. Secondary growth is supported by the lateral meristem or vascular cambium, whose division and differentiation form the secondary xylem towards the inside of the stem and form the secondary phloem towards the outside of the stem (Du and Groover, 2010). After differentiation, SCW is thickened and lignified to form wood.

In plants, SCW biosynthesis is regulated by a hierarchical NAC-MYB gene regulatory network, in which the plant-specific NAC (NAM, ATAF, and CUC) transcription factors (TFs) act as master regulators that directly target MYB46/83 TFs, which switch on biosynthesis of all three major SCW components (lignin, cellulose and xylan) by directly activating the expression of a number of MYB TFs and SCW biosynthetic genes (Xie et al., 2018a). For example, Arabidopsis SECONDARY WALL-ASSOCIATED NAC DOMAIN PROTEIN 1/NAC SECONDARY WALL THICKENING PROMOTING FACTOR 3 (SND1/NST3) and its close homologs NST1, NST2, VASCULAR-RELATED NAC DOMAIN6 (VND6) and VND7 were found to form the first layer of the NAC-MYB gene regulatory network and regulates SCW formation in various tissues (Xie et al., 2018a). In poplar, a group of NAC TFs named WOOD-ASSOCIATED NAC DOMAIN PROTEINs (WNDs) were found to be functional orthologs of Arabidopsis NAC TFs (Zhong et al., 2010; Zhong and Ye, 2010). In particular, PtrWND2B and PtrWND6B have been shown to directly activate the expression of poplar MYB46/83 homologs, many other transcription factors and synthetic genes involved in SCW biosynthesis (Zhong et al., 2010). Dominant suppression of PtrWND2B or PtrWND6B in poplar reduced cell wall thickness of xylem fibers by over 50% (Zhong et al., 2010).

SCW biosynthesis and its transcriptional regulation in poplar are highly plastic in response to environmental signals (Zinkgraf et al., 2017). The expression of cell wall-associated genes is tightly spatiotemporally co-regulated (Brown et al., 2005). Environmental stresses such as drought and salt have been shown to reduce the total amount of cell wall polysaccharides, vessel size and S/G lignin ratio of poplar stems (Hori et al., 2020). Iron (Fe) is a micronutrient that is essential for plant redox processes and photosynthesis (Briat et al., 2015). Fe deprivation was found to increase SCW deposition and lignification in root via a REVOLUTA (REV)-controlled gene regulatory network (Taylor-Teeples et al., 2015). However, how Fe deprivation affects SCW formation in aboveground tissues, which are the main source of biomass, is still poorly studied.

The basic helix-loop-helix (bHLH) TF family is one of the largest TF families in plants. The bHLH TFs have been found to play vital roles in plant growth, SCW biosynthesis, iron homeostasis and stress responses (Gao and Dubos, 2023). Phylogenetic studies divided the family into 17 major groups (I–XVII) and 31 subfamilies (Pires and Dolan, 2010; Gao and Dubos, 2023). Among them, members of the bHLH IVb subgroup have been reported to regulate iron homeostasis through various mechanisms. AtbHLH011, UPSTREAM REGULATOR OF IRT1 (URI/AtbHLH121) and POPEYE (PYE/AtbHLH047) are three bHLH IVb TFs in Arabidopsis. AtbHLH11 functions as a negative regulator of FER-LIKE IRON DEFICIENCY-INDUCED TRANSCRIPTION FACTOR (FIT)-dependent Fe uptake (Tanabe et al., 2019). AtbHLH11 has no transcriptional activity but interacts with bHLH IVc TFs to repress their transactivity on bHLH Ib TFs (Tanabe et al., 2019). URI/bHLH121 is expressed throughout the plant body and acts as a direct transcriptional activator of key genes involved in the Fe regulatory network (Gao et al., 2020). URI is phosphorylated upon Fe deficiency, forms heterodimers with bHLH IVc TFs, and induces transcription of bHLH Ib TFs, which in turn heterodimerize with FIT and drive transcription of IRON-REGULATED TRANSPORTER1 (IRT1) and FERRIC REDUCTION OXIDASE2 (FRO2) to increase Fe uptake (Kim et al., 2019). Shoot iron deficiency signal activates transcription of *PYE*/*AtbHLH047*, which encodes a mobile protein to facilitate iron uptake, root-to-shoot translocation, and storage via FIT-dependent and/or - independent pathways (Long et al., 2010; Muhammad et al., 2022).

In this study, we found that Fe deprivation is capable of activating SCW biosynthesis via its gene regulatory network in poplar stem. A bHLH IVb TF (PtrbHLH011) in poplar, the close homolog of AtbHLH011, appears to be involved in this regulation. *PtrbHLH011* expression is downregulated during wood formation in stems and by iron deprivation in various plant tissues. Overexpression of *PtrbHLH011* in poplar inhibited growth and reduced stem SCW formation. In leaf tissue, flavonoid reduction, iron and zinc disorders, and necrotic leaf spots were found to be induced by *PtrbHLH011* overexpression. Our studies integrating RNA-seq, transient ChIP-seq and transactivation assay revealed that PtrbHLH011 is a transcriptional repressor and plays diverse regulatory roles in SCW biosynthesis, flavonoid biosynthesis, and metal homeostasis by directly repressing the expression of key regulators and genes for these biological pathways. We found that PtrbHLH011 targets these genes by directly binding to the *cis*-regulatory element with an AAAGACA sequence. In summary, our studies demonstrated that PtrbHLH011 has multifaced functions in SCW biosynthesis, flavonoid biosynthesis, and metal homeostasis, and may act as a co-regulator for coordinating wood formation and systematic responses to Fe deprivation in poplar.

## Results

### Fe deprivation activates SCW biosynthesis

We found that Fe-deprivation stress significantly increases lignin and SCW deposition in poplar stems. The Fe-deprivation treatment was performed by growing poplar seedlings in a hydroponic nutrient solution without iron (0 μM Fe) for 21 days (Methods and Fig. S1). In parallel, poplar seedlings grown in a hydroponic nutrient solution with sufficient iron for poplar (10 μM Fe) was used as a control. As previously reported (Chen et al., 2019), our Fe-deprivation treatment induced leaf chlorosis and stem growth inhibition (Fig. S1). Stem cross-sections from 3^rd^ and 10^th^ internodes were then collected for lignin staining with phloroglucinol-HCl. As shown in Fig. 1A, B, S2, ectopic lignin and SCW deposition was observed in developing xylem and phloem of the stems with primary growth (3^rd^ internode). Similarly, in the stems with secondary growth (10^th^ internode), over 100% increase in the SCW thickness of xylem fibers was observed under Fe deprivation compared to the control (Fig. 1C-E and Fig. S2).

**Figure 1.**
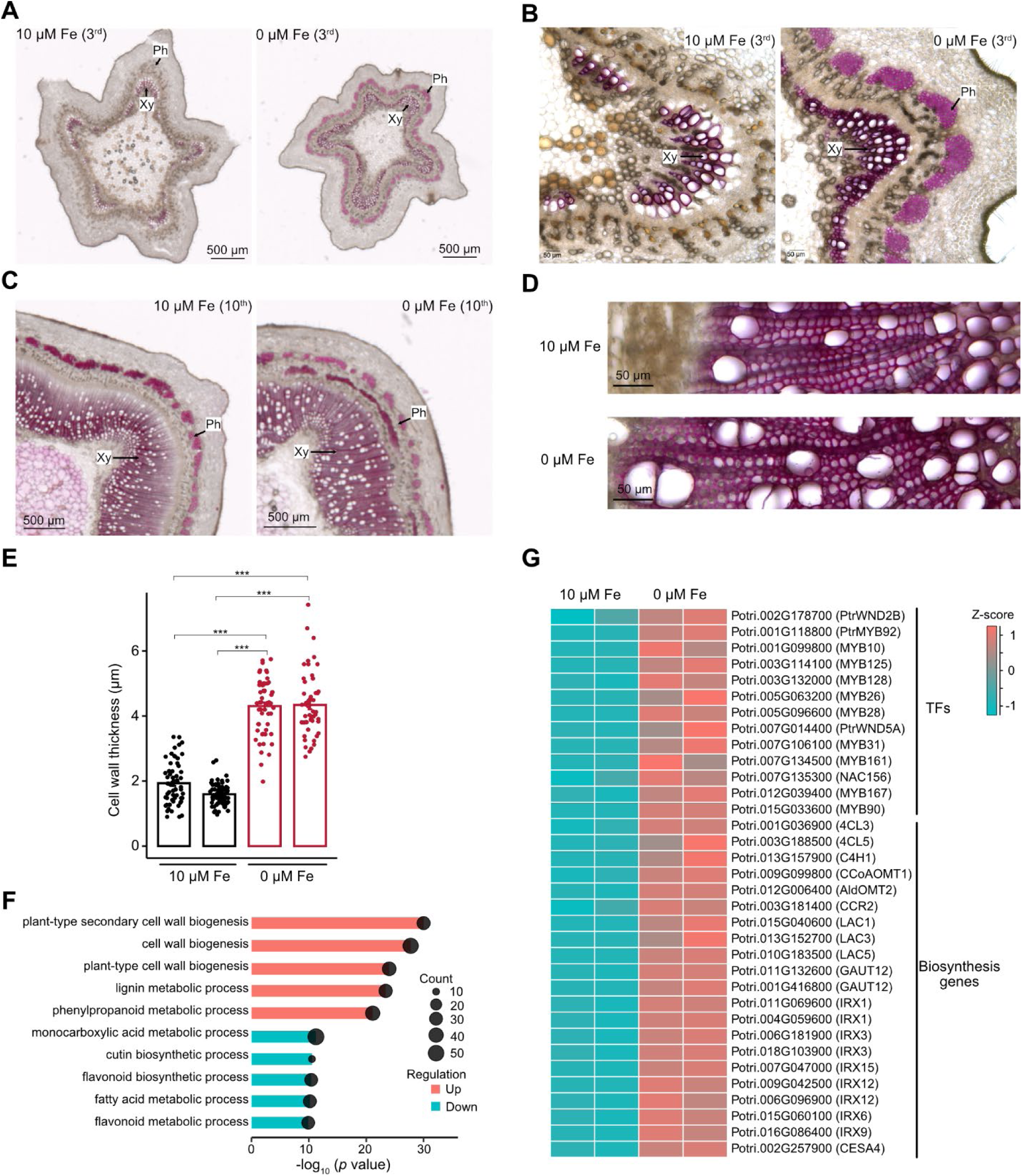
Iron (Fe) deprivation activates secondary cell wall (SCW) biosynthesis in poplar stems. (**A** and **B**) images of phloroglucinol-HCl stained cross-sections of the 3^rd^ internode of poplar treated with control (10 μM Fe) and Fe deprivation (0 μM Fe). **(C** and **D)** images of phloroglucinol-HCl stained cross-sections of the 10^th^ internode of poplar treated with control (10 μM Fe) and Fe deprivation (0 μM Fe). Xy, of xylem fiber cells by Fe deprivation. Two biological replicates of control (black) and two biological replicates of Fe deprivation (red) treatment are shown. Student’s *t*-tests were performed. ***, *P* < 0.001. (**F**) bar plot showing the top five enriched gene ontology (GO) terms among upregulated genes (red) and downregulated genes (blue) by Fe deprivation in 3^rd^ internode. (**G**) heatmap showing Fe deprivation-induced upregulation of transcription factors (TFs) and biosynthesis genes essential for SCW biosynthesis in the 3^rd^ internode. RNA-seq data of two replicates for control and Fe deprivation conditions are shown.

Consistently, RNA-seq analysis revealed that Fe deprivation activates the expression of various transcriptional activators and synthetic genes for SCW biosynthesis (Fig. 1F, G). The 3^rd^ internode had the most altered transcriptomes by Fe deprivation with 1,211 upregulated and 1,038 downregulated genes compared to the control (Fig. S3 and Data S1, S2). Among the upregulated genes, all the top five enriched gene ontology (GO) terms are relevant to lignin and SCW biosynthesis (Fig. 1F). Furthermore, a large set of TFs and biosynthesis genes for SCW were upregulated by Fe deprivation (Fig. 1G). Taken together, our results suggest that Fe deprivation activates SCW biosynthesis through transcriptome reprogramming.

### PtrbHLH011 is involved in Fe deprivation-induced SCW biosynthesis

Since transcriptome reprogramming of Fe homeostasis is also a key mechanism in response to Fe deprivation (Buckhout et al., 2009), we hypothesized that transcriptional co-regulators may exist to control SCW biosynthesis and Fe homeostasis pathways more cost-effectively. bHLH transcription factors play essential roles in regulating plant Fe homeostasis (Gao and Dubos, 2023). By searching AspWood, a high-spatial-resolution RNA-seq database across developing phloem and wood-forming tissues in poplar (Sundell et al., 2017), for bHLH transcription factors that may also regulate SCW biosynthesis, we identified *Potri.005G113400*. We named this gene PtrbHLH011 because it encodes a bHLH IVb TF that is a close homolog of AtbHLH011 (Fig. 2A and Fig. S4). Across the four replicates in AspWood, *PtrbHLH011* exhibits a consistent expression pattern in wood-forming tissues that appears to be opposite to the expression patterns of SCW biosynthesis marker genes *PtrWND2B* and *PtCesA4* (Zhong et al., 2010; Sundell et al., 2017), suggesting that PtrbHLH011 may negatively regulate SCW biosynthesis (Fig. 2B). Similar to its Arabidopsis homolog AtbHLH011, a negative regulator of Fe uptake, *PtrbHLH011* expression is significantly downregulated by Fe-deprivation treatment. qRT-PCR results showed that *PtrbHLH011* expression decreased by over 50% under Fe-deprivation treatment in tissues including xylem, phloem, leaf and root (Fig. 2C).

**Figure 2.**
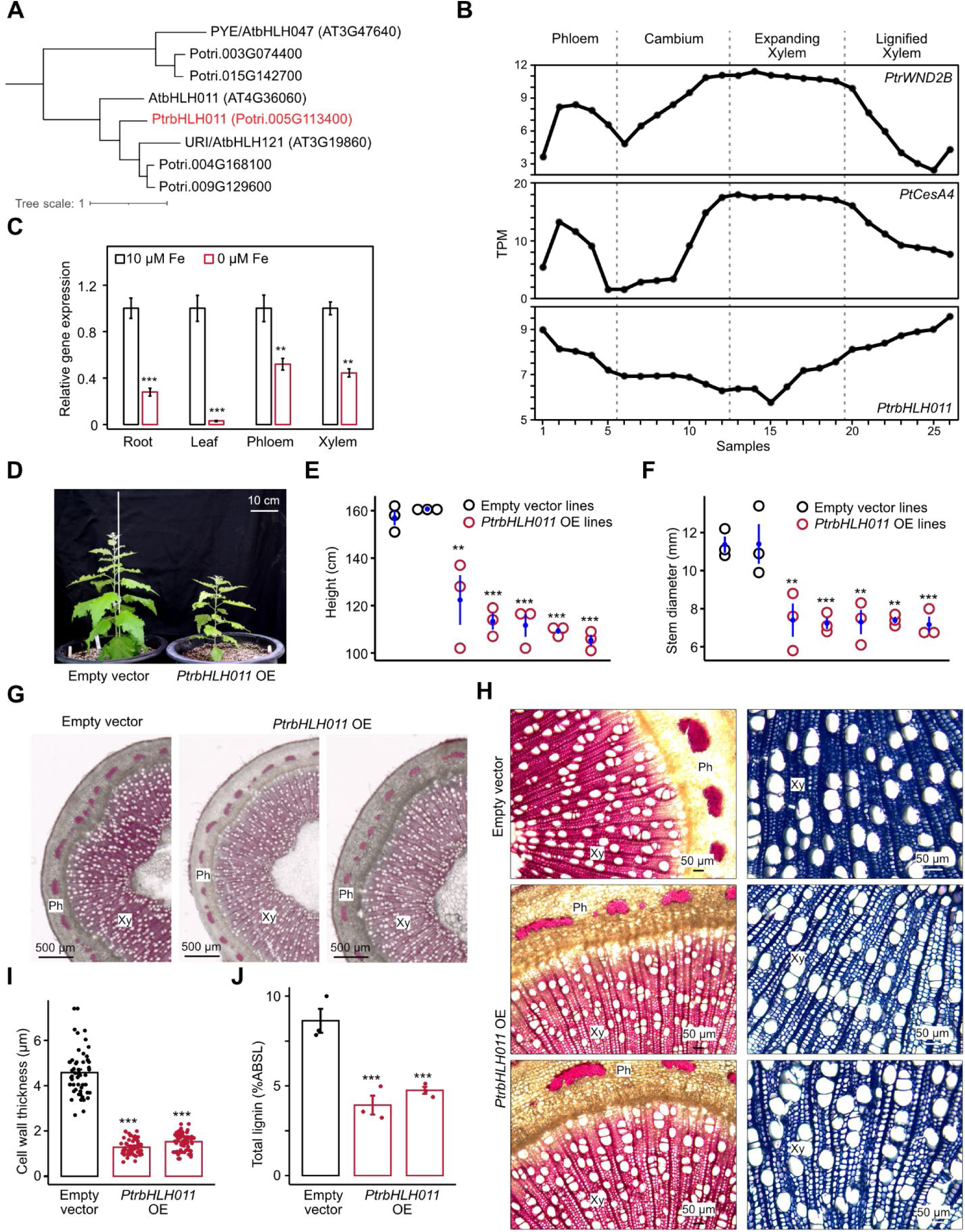
PtrbHLH011 inhibits secondary cell wall (SCW) biosynthesis. (**A**) Phylogenetic clade highlighted by red color. Bootstrap values are 92 and above based on an ultrafast bootstrap analysis. The full tree is available in **Fig. S4**. (**B**) Expression patterns of *PtrWND2B*, *PtCesA4* and *PtrbHLH011* in wood-forming tissues. The data were retrieved from Aspwood database (https://plantgenie.org). The three genes have consistent expression patterns in the four replicates and the normalized RNA-seq counts (TPM, transcript per million) of one representative replicate are shown. Vertical dashed lines indicate transcriptome reprogramming events (left to right): phloem and xylem differentiation, onset of SCW formation, and end of SCW deposition. (**C**) qRT-PCR results showing the expression of *PtrbHLH011* is downregulated by iron (Fe) deprivation (0 μM Fe) treatment in root, leaf, phloem and xylem. Gene expression was normalized against poplar *eIF-5A* gene, with control (10 μM Fe) set as 1. Values represent means ± SE, n=3. Two-tailed Student’s *t*-tests were performed by comparing with control. (**D**) Image comparing two-month-old transgenic poplar plants with empty vector and *PtrbHLH011* overexpression (*PtrbHLH011* OE). (**E** and **F**) Dot plots comparing plant height and stem diameter of four-month-old transgenic poplar plants with empty vector and *PtrbHLH011* OE. Three biological replicates of two independent empty vector lines (black) and five independent *PtrbHLH011* OE lines (red) were measured. The mean ± SE of three biological replicates is shown in blue color. Two-tailed Student’s *t*-tests were performed by comparing each *PtrbHLH011* OE line with the six empty vector plants. (**G** and **H**) Phloroglucinol-HCl and toluidine blue staining of cross-sections of the 10^th^ internode demonstrating the reduction of lignin and SCW in *PtrbHLH011* OE lines. One empty vector line and two representative *PtrbHLH011* OE lines are shown. Xy, developing xylem; Ph, developing phloem. (**I**) Bar plot showing the reduced cell wall thickness of xylem fiber cells in *PtrbHLH011* OE lines. (**J**) Reduced total lignin content in stems of *PtrbHLH011* OE lines. %ABSL, percentage of acetyl bromide soluble lignin. Student’s *t*-tests were performed. In **C, E, F, I, J**, **, *P*<0.01; ***, *P*<0.001.

Additionally, we observed significant reductions in stem SCW in transgenic poplars overexpressing *PtrbHLH011*, which is consistent with our hypothesis. As shown in Fig. 2D-F and Fig. S5A, all five independent *PtrbHLH011* overexpression (*PtrbHLH011* OE) lines showed significantly inhibited growth compared to the empty vector control, with a 29.2% reduction in plant height (Fig. 2E) and a 35.9% reduction in stem diameter on average (Fig. 2F). By phloroglucinol-HCl and toluidine blue staining of stem cross-sections of two representative *PtrbHLH011* OE lines, we observed a reduction in SCW thickness of xylem and phloem cells by over 65% (Fig. 2G-I and Fig. S5B). Furthermore, we detected an over 45% reduction in stem lignin content in *PtrbHLH011* OE plants (Fig. 2J). Taking together, our results indicate that PtrbHLH011 inhibits SCW and lignin biosynthesis.

### PtrbHLH011 affects flavonoid biosynthesis and iron homeostasis

The *PtrbHLH011* OE lines also have a smaller leaf size than the empty vector control (Fig. 3A). In addition, leaf chlorosis and necrotic leaf spots were observed on *PtrbHLH011* OE leaves (Fig. 3A). To investigate the causes of these disease symptoms, we first measured the accumulation of flavonoids, the main defense metabolites produced through the phenylpropanoid pathway (Fini et al., 2011). As shown in Fig. 3B and Fig. S6, the soluble phenolic and anthocyanin contents in the leaves of *PtrbHLH011* OE lines showed a significant reduction. Anthocyanins are a class of flavonoids responsible for leaf color and stress tolerance (Li and Ahammed, 2023). Its reduction likely results in foliar symptoms of *PtrbHLH011* OE plants.

**Figure 3.**
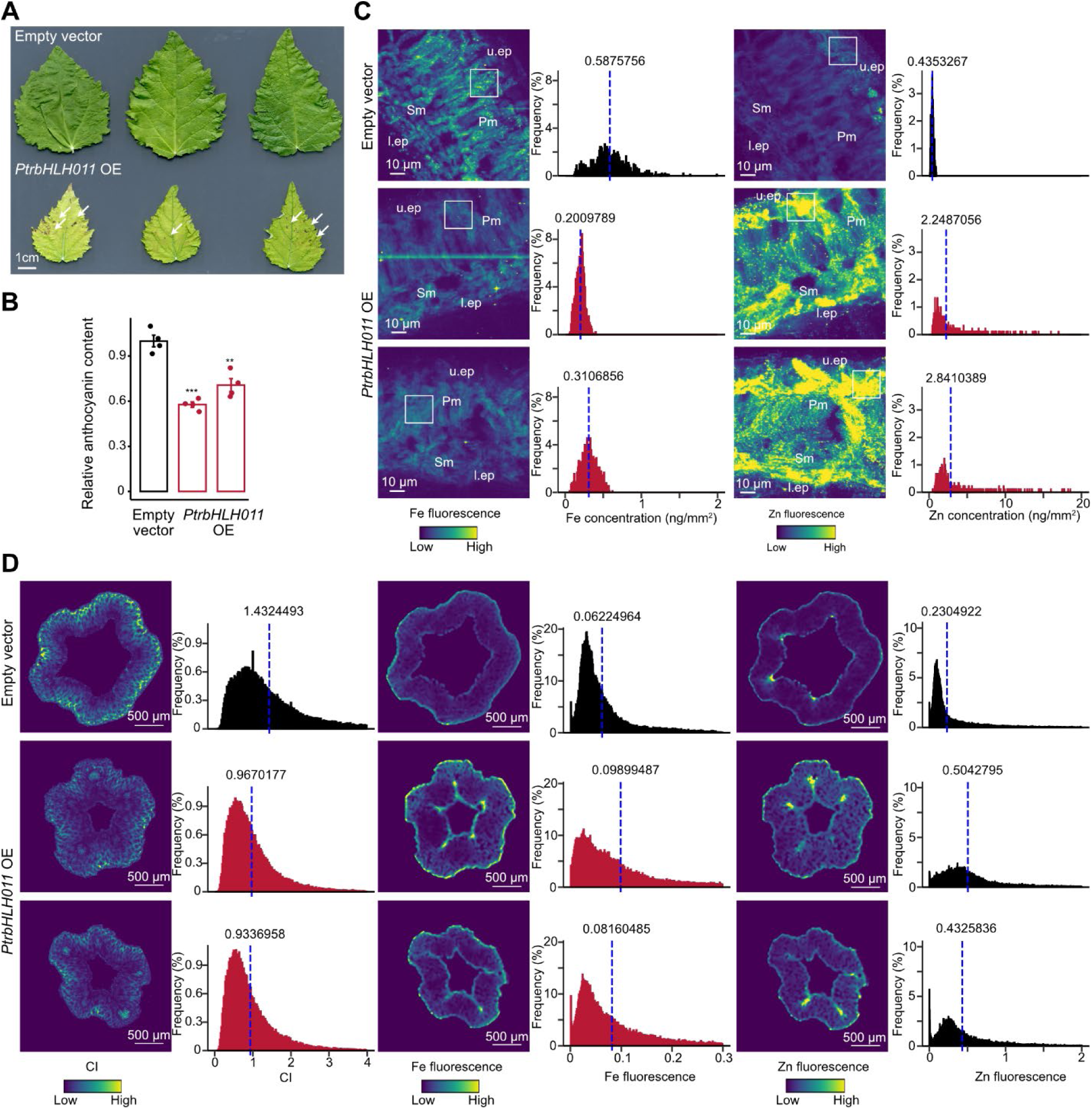
PtrbHLH011 affects flavonoid biosynthesis and metal homeostasis. (**A**) Image comparing young leaves of empty vector and *PtrbHLH011* overexpression (*PtrbHLH011* OE) transgenic poplar. White arrows indicate necrotic leaf spots. (**B**) Bar plot showing reduced anthocyanin content in two representative *PtrbHLH011* OE lines. Two-tailed Student’s *t*-tests were performed by comparing with empty vector control. **, *P*<0.01; ***, *P*<0.001. (**C**) X-ray fluorescence microscopy of leaf cross-sections showing reduced iron (Fe) and increased zinc (Zn) content in leaf cells of *PtrbHLH011* OE plants. Histograms on the right of the images show the distribution of fluorescence signals in the selected spongy mesophyll; u.ep, upper epidermis; l.ep, lower epidermis. (**D**) 2-D reconstructed images of the peeled stem cross-sections of empty vector control and two representative *PtrbHLH011* OE plants. Images were reconstructed using X-ray scattering data (cellulose crystallinity index, CI) and X-ray fluorescence data (Fe and Zn). Histograms on the right of the images show the distribution of scattering or fluorescence signals in reconstructed images. The mean value is indicated by a blue dashed line.

Furthermore, by performing X-ray fluorescence microscopy (XRF) analysis, we discovered altered Fe accumulation in *PtrbHLH011* OE plants. We found that leaves from *PtrbHLH011* OE lines have reduced Fe content in the palisade mesophyll layer (Fig. 3C), where the majority of photosynthesis occurs. In general, Zn and Fe compete for the same transporter, leading to opposite accumulation patterns under Fe-deprivation condition(Baxter et al., 2008). Consistently, in *PtrbHLH011* OE leaves, we observed increased zinc (Zn) content in the epidermal layers (Fig. 3C), where most Zn accumulates. Due to the importance of Fe in the chloroplast and the toxicity of high Zn to plant tissues, their disorders in *PtrbHLH011* OE leaves may also be a cause of the leaf symptoms. Additionally, we performed X-ray scattering and fluorescence analysis on mature stems (14^th^ internode) of the same plants. As expected, the X-ray scattering results clearly showed the changes in SCW in the xylem of *PtrbHLH011* OE plants, as indicated by the reduction in cellulose crystallinity index (CI; Fig. 3D). The X-ray fluorescence results showed that the Fe and Zn contents in the xylem of *PtrbHLH011* OE stems have a moderate increase compared to the empty vector control, suggesting that the translocation of Fe and Zn towards leaf tissues may be impaired.

### PtrbHLH011 is a transcriptional repressor that directly represses key genes for SCW biosynthesis, flavonoid biosynthesis, and metal homeostasis

The PtrbHLH011 protein is localized in both the cytosol and the nucleus in poplar mesophyll protoplasts (Fig. 4A). Furthermore, our protoplast-based transient transactivation assays using the GUS and luciferase reporter system (Xie et al., 2018b) revealed that PtrbHLH011 has transcriptional repressor activity but not activator activity (Fig. 4B, C). PtrbHLH011 was fused with the Gal4 DNA-binding domain (GD-PtrbHLH011) and recruited to the promoter of the GUS reporter gene via the interaction between GD and the Gal4 DNA sequence upstream of GUS. A plasmid overexpressing luciferase (LUC) was co-transfected as an internal standard. To test repressor activity, an additional transactivator plasmid overexpressing LexA DNA-binding domain (LD) fused herpes simplex virus VP16 transactivator (VP16) was co-transfected to constitutively activate GUS expression through the interaction between LexA and LD. The normalized GUS activity (GUS/LUC) was used to evaluate transcriptional activity of PtrbHLH011. As shown in Fig. 4B, PtrbHLH011 has a significantly lower GUS/LUC value than the green fluorescent protein (GFP) negative control, indicating transcriptional repressor activity. In contrast, the GUS/LUC value of PtrbHLH011 is similar to that of the GFP negative control and much less than that of the VP16 positive control in the activation assay (Fig. 4C), suggesting that PtrbHLH011 does not have transcriptional activator activity.

**Figure 4.**
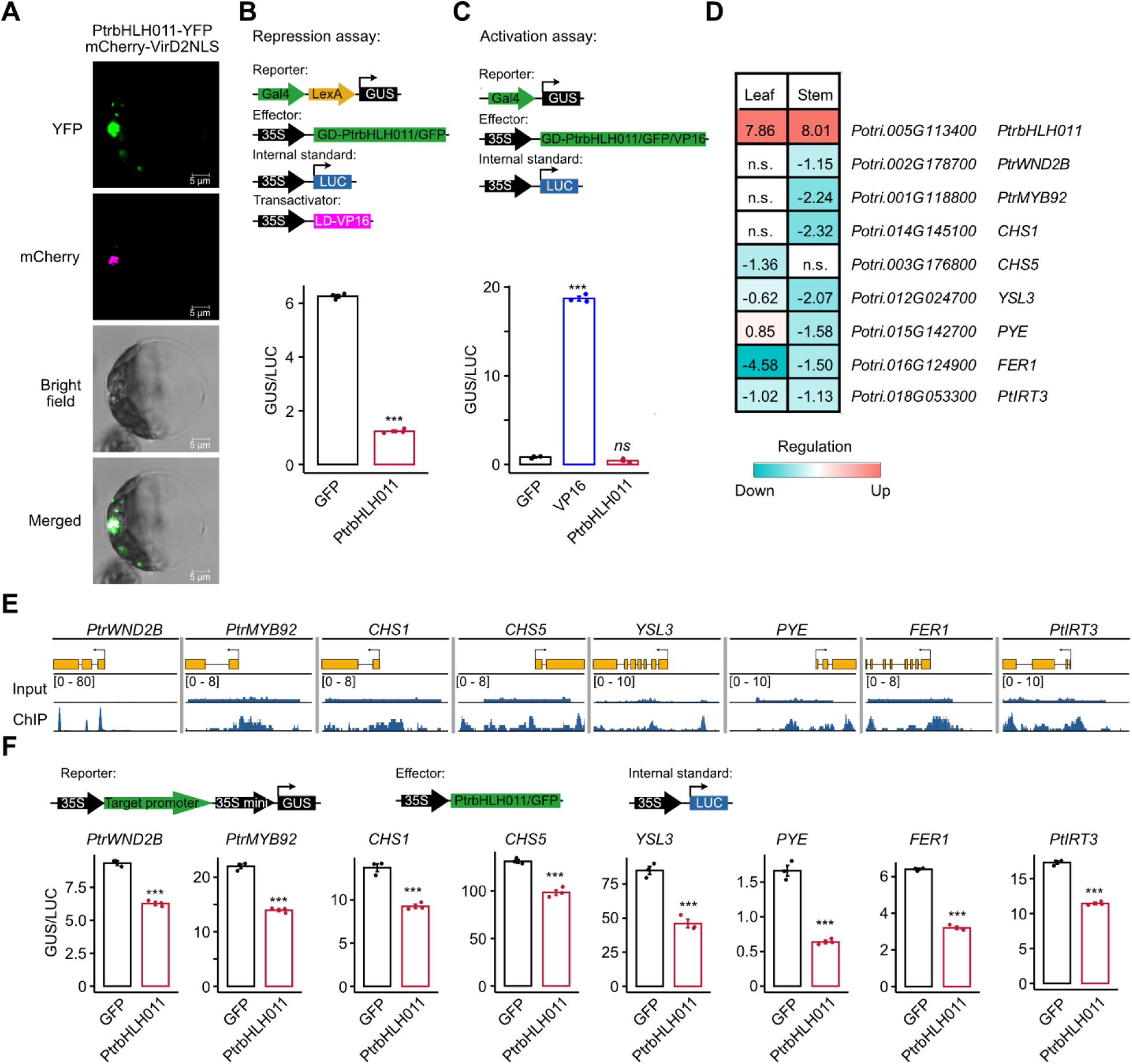
PtrbHLH011 is a transcriptional repressor that directly targets key genes for secondary cell wall (SCW) biosynthesis, flavonoid biosynthesis, and metal homeostasis. (**A**) Subcellular localization of PtrbHLH011 in poplar protoplasts. Yellow fluorescent protein tagged PtrbHLH011 (PtrbHLH011-YFP; green) and the nuclear marker (mCherry-VirD2NLS; magenta) were coexpressed in protoplasts and their fluorescent signal overlap is indicated in white. (**B**) Transient transactivation assay for PtrbHLH011 repressor activity. The top schematic displays the four vector types used. The transactivator construct overexpressing VP16 fused with the LexA DNA-binding domain (LD) was used to constitutively activate GUS expression through the interaction between LexA and LD. Green fluorescent protein (GFP) was used as a negative control. (**C**) Transient transactivation assay for negative control and VP16 was used as a positive control. (**D**) Heatmap showing the reduction in gene expression of key genes for SCW biosynthesis, flavonoid biosynthesis and iron homeostasis in leaf and/or stem tissue of transgenic poplar with *PtrbHLH011* overexpression (*PtrbHLH011* OE). Numbers in the table are log_2_ (fold change) values in RNA-seq analysis. n.s., not significant compared to the empty vector control (*P* ≥ 0.05). (**E**) Transient ChIP-seq read depths in *WND2B*, *MYB92*, *CHS1*, *CHS5*, *YSL3*, *PYE*, *FER1* and *ZIP4* loci. Read depths of three biological replicates mapped to the *Populus alba* genome were averaged and shown. Read depths mapped to the *Populus tremula* genome have similar patterns. Gene coding regions and 2-kb promoter regions are shown. (**F**) Transient transactivation assays show that PtrbHLH011 targets and represses *WND2B*, *MYB92*, *CHS1*, *CHS5*, *YSL3*, *PYE*, *FER1* and *ZIP4*. The top schematic displays the three vector types used. Relative GUS activity (GUS/LUC) was calculated by normalizing GUS activity against luciferase activity (**B, C, F**). Values represent means + SE, n=3 (**B, C, F**). *P* values were calculated by Student’s *t*-tests (**B, C, F**). ***, *P*<0.001; *ns*, *P*≥0.05.

To seek genome-wide targets of PtrbHLH011 repression, we integrated RNA-seq and protoplast-based transient ChIP-seq approaches (Tadesse et al., 2024). By comparing the transcriptomes of *PtrbHLH011* OE and empty vector control, we identified 2,743 upregulated and 1,227 downregulated genes in the leaf tissue, 2,172 upregulated and 1,854 downregulated genes in the stem tissue (Fig. S7A-D and Data S3, S4). In transient ChIP-seq experiments, PtrbHLH011 exhibited the typical binding peak of transcription factors surrounding the transcription start site (TSS; Fig. S7E, F). By identifying downregulated genes in either tissue whose promoters have PtrbHLH011 binding peaks (Fig. 4D,E and Data S5), we discovered putative PtrbHLH011 targets that are responsible for SCW biosynthesis, flavonoid biosynthesis and iron homeostasis, including *PtrWND2B* and *PtrMYB92*, which are major activators of SCW biosynthesis (Zhong et al., 2010; Liu et al., 2021); *CHALCONE SYNTHASE 1* and *5* (*CHS1* and *CHS5*), which encode the enzyme that catalyzes the first step of flavonoid biosynthesis; YELLOW STRIPE LIKE 3 (YSL3) and PYE, the Arabidopsis homologs of which are responsible for root-to-shoot iron translocation and long-distance iron signaling (DiDonato Jr et al., 2004; Long et al., 2010; Kumar et al., 2017; Muhammad et al., 2022); and IRON REGULATED TRANSPORTER 3 (PtIRT3) and FERRITIN 1 (FER1) for shoot iron accumulation (Lin et al., 2009; Huang and Dai, 2015).

Noting significant PtrbHLH011 binding peaks in the promoters of these genes (Fig. 4E), reporter constructs were generated by inserting the *PtrWND2B*, *PtrMYB92*, *CHS1*, *CHS5*, *YSL3*, *PYE*, *PER1* and *PtIRT3* promoters between the 35S promoter and the GUS reporter gene, respectively (Fig. 4F). The transient transactivation assay confirmed that PtrbHLH011 directly represses the activity of all these promoters (Fig. 4F).

### PtrbHLH011 binds to the AAAGACA motif and represses the expression of target genes in plant tissues

The repression of PtrbHLH011 on these eight target genes was further validated in *PtrbHLH011* OE plants. qRT-PCR results show that all eight genes except *PYE* are downregulated in leaf and stem tissues (Fig. 5A). *PYE* had a significant reduction in the stem, but not in the leaf (Fig. 5A). This could be due to the major function of PYE in iron translocation in root and stem.

**Figure 5.**
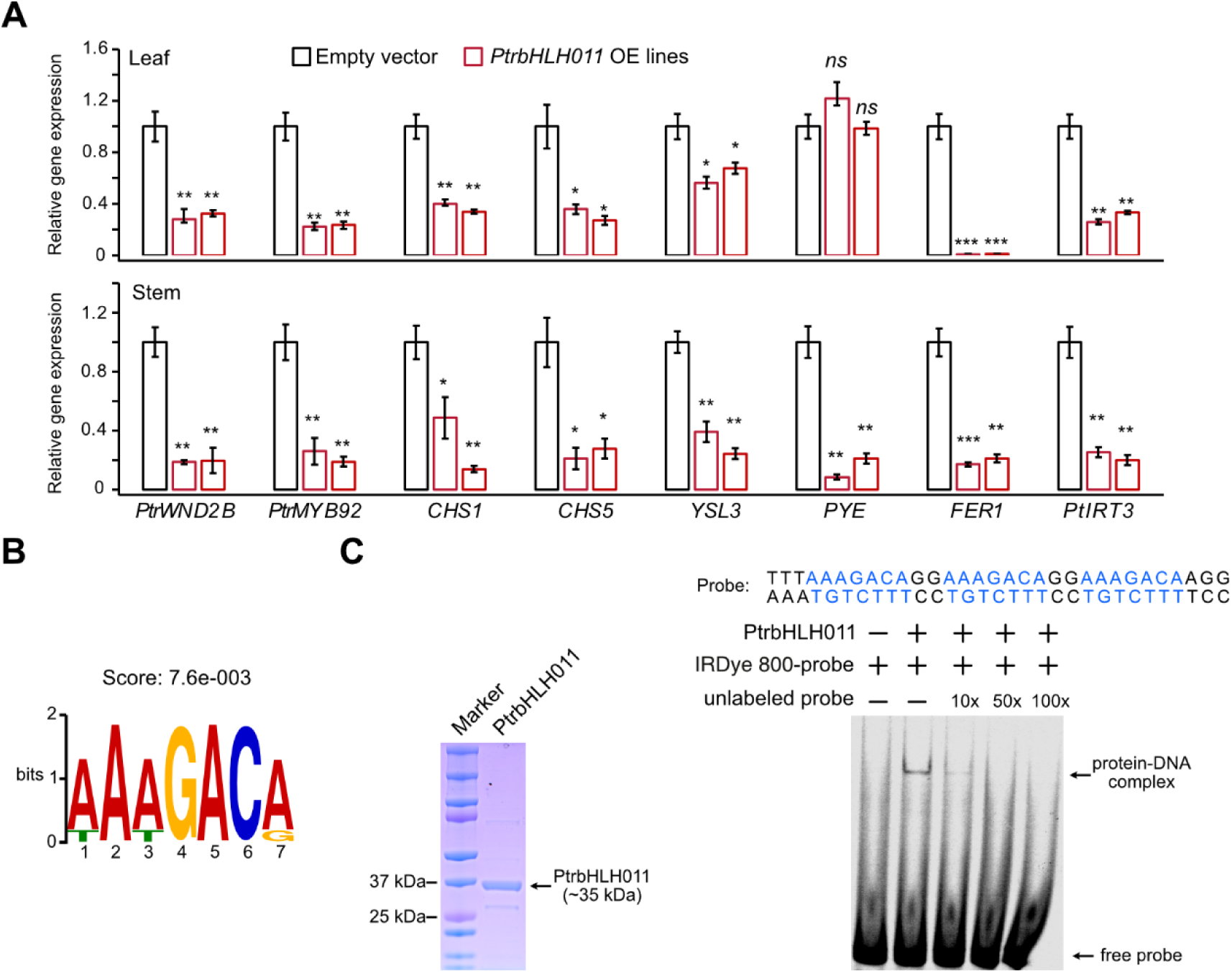
PtrbHLH011 binds to the *cis*-element AAAGACA to repress the expression of target genes. (**A**) qRT-PCR results showing the expression of PtrbHLH011 targets in root, leaf, and stem tissues of an empty vector line and two representative *PtrbHLH011* OE lines. Gene expression was normalized against poplar *eIF-5A* gene, with empty vector set as 1. Values represent means ± SE, n=3. Student’s *t*-tests were performed by comparing with control. ***, *P*<0.001; **, *P*<0.01; *, *P*<0.05; *ns*, *P*≥0.05. (**B**) The AAAGACA motif is enriched in promoters of validated PtrbHLH011 targets. (**C**) PtrbHLH011 binds to the AAAGACA motif *in vitro*. Left: Coomassie blue stained protein gel showing the purified PtrbHLH011 protein. Right: Electrophoretic Mobility Shift Assay (EMSA) showing that PtrbHLH011 specifically binds to the DNA probe containing three copies of the AAAGACA sequence (blue color in the probe sequence).

By investigating *cis*-regulatory elements enriched in promoters of the eight PtrbHLH011 targets, but not in promoters of genes not directly repressed by PtrbHLH011 in the transactivation assay (Fig. S8A and Methods), we identified the sequence AAAGACA (Score: 7.6e-003) as a potential *cis*-regulatory element bound by PtrbHLH011 (Fig. 5B). The promoters of the eight PtrbHLH011 targets contain at least one copy of this AAAGACA motif (Fig. S8B).

Electrophoretic mobility shift assay (EMSA) confirmed that PtrbHLH011 specifically binds to this DNA motif (Fig. 5C). We expressed and purified the full-length PtrbHLH011 protein (Fig. 5C) and synthesized 5’ IRDye 800-labeled DNA probes containing three copies of the AAAGACA sequence (Fig. 5C). EMSA and the competition assay using 10x, 50x and 100x unlabeled probe of the same sequence showed that the binding was specific, as indicated by the reduction of the bound signal (protein-DNA complex) in the presence of the competitor (Fig. 5C).

## Discussion

Growing bioenergy crops on marginal lands with low nutrient quality is increasingly utilized as a solution to minimize competition for arable land and food production (Gelfand et al., 2013). However, as the major source of lignocellulosic biomass, SCW biosynthesis is highly plastic in response to environmental stress (Zinkgraf et al., 2017). For the massive production of lignocellulosic biomass on marginal lands that are often exposed to various abiotic stresses, it is crucial to understand how these stresses affect the quantity and quality of SCW. Additionally, the discrepancies between field-grown and greenhouse-grown cell-wall-engineered poplars are increasingly observed (De Meester et al., 2022; Li et al., 2023), which is one of the main reasons for hampering the commercialization of wood feedstocks for biofuels worldwide. The lack of knowledge about the adaptation of SCW biosynthesis to environmental stress is a major factor underlying the discrepancies.

Iron is an essential micronutrient for photosynthesis, respiration and other important developmental processes in plants (Briat et al., 2015). However, iron deficiency caused by poor solubility and alkaline soils is widespread and detrimental to crop yield and bioproduct quality. To date, Fe deprivation has only been reported to increase SCW deposition and lignification of root in the model plant Arabidopsis via the REV-controlled gene regulatory network (Taylor-Teeples et al., 2015). In this study, we demonstrated that iron deprivation upregulates the gene regulatory network of SCW and increases SCW biosynthesis in poplar stems, the main source of lignocellulosic biomass. We identified a bHLH IVb transcription factor PtrbHLH011 that appears to be involved in this regulation. We found that PtrbHLH011 is a potent inhibitor of SCW biosynthesis by directly targeting PtrWND2B and PtrMYB92, two major activators of SCW biosynthesis that were found to activate the expression of numerous transcription factors and biosynthesis genes for SCW biosynthesis in poplar (Zhong et al., 2010; Liu et al., 2021). Under iron deficiency, *PtrbHLH011* expression is downregulated to relieve the repression of *PtrWND2B* and *PtrMYB92*, which consequently activates SCW biosynthesis. Therefore, we propose that PtrbHLH011 acts as a gatekeeper by limiting SCW biosynthesis under normal growth conditions to prevent growth inhibition caused by excessive lignification (Fig. 6). Fe deficiency signal silences *PtrbHLH011* expression to enhance lignification of vascular tissue for better water and iron transportation from root to shoot (Fig. 6). However, it does not exclude the involvement of other transcription factors. Screening additional co-regulators and knockout study of *PtrbHLH011* could further elucidate the molecular mechanisms and GRNs controlling SCW biosynthesis under Fe deficiency.

**Figure 6.**
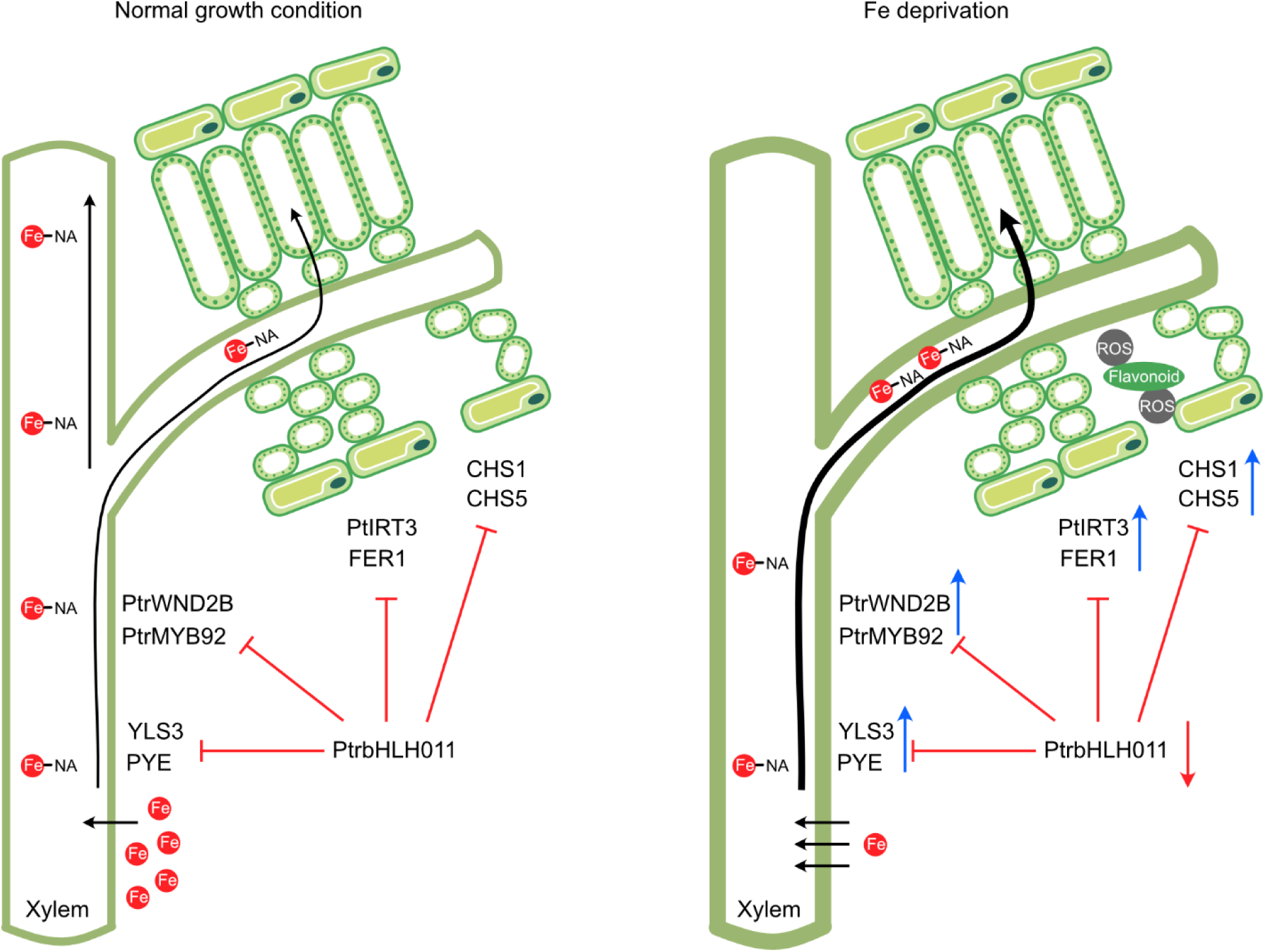
Proposed model of the central regulatory role of PtrbHLH011 in response to iron (Fe) deprivation. PtrbHLH011 directly represses the expression of PtrWND2B and PtrMYB92 for secondary cell wall (SCW) biosynthesis, CHS1 and CHS5 for flavonoid biosynthesis, YSL3 and PYE for root-to-shoot iron transport and Fe deficiency signaling, as well as PtIRT3 and FER1 for shoot Fe accumulation. Under normal growth conditions (left), PtrbHLH011 acts as a gatekeeper to prevent excessive lignification, secondary metabolism and Fe accumulation, all of which inhibit plant growth. Under Fe deprivation (right), PtrbHLH011 is downregulated to release these repressions to promote xylem lignification for iron and water transport, to activate Fe deficiency signaling and iron homeostasis mechanisms for shoot Fe transport and accumulation, and to enhance flavonoid biosynthesis for the oxidative stress caused by Fe deficiency. Black arrows indicate Fe transport, red T shapes indicate transcriptional repression, red arrow indicates downregulation, and blue arrows indicate upregulation.

We showed that PtrbHLH011 is a crucial regulatory node for Fe homeostasis in poplar. Fe is an essential micronutrient for plant growth and development. However, excessive amounts of Fe delivered to cells can be harmful due to the redox properties of Fe (Marschner, 2011). We found that PtrbHLH011 directly represses the expression of key genes for Fe deficiency signaling, Fe translocation and shoot Fe accumulation, including *PYE*, *YSL3, PtIRT3* and *FER1*. PYE is a Fe deprivation-induced bHLH TF and was reported to play an important role in the sense of shoot Fe deprivation signal to increase Fe uptake from soil and root-to-shoot Fe translocation in Arabidopsis (Long et al., 2010; Muhammad et al., 2022). In dicots, YSL proteins are responsible for the mobilization of nicotianamine (NA)-chelated Fe in the vasculature (DiDonato Jr et al., 2004; Waters et al., 2006). Arabidopsis homologs of YSL3, AtYSL1 and AtYSL3, have been reported to be required for long-distance Fe deficiency signaling from shoot to root (Kumar et al., 2017). PtIRT3 is an Fe transporter that is constitutively expressed in several plant tissues (Huang and Dai, 2015). Its Arabidopsis homolog IRT3 also plays an essential role in maintaining shoot Fe accumulation (Shanmugam et al., 2011). Additionally, overexpression of *FER* genes have been reported to increase leaf Fe accumulation (Van Wuytswinkel et al., 1999; Goto et al., 2000). To prevent excessive Fe accumulation in leaf cells, PtrbHLH011 limits the expression of *YSL3*, *PYE*, *PtIRT3* and *FER1* under normal growth conditions. This may result in reduced Fe accumulation in leaf cells and increased Fe accumulation in xylem in *PtrbHLH011* OE plants (Fig. 3). On the other hand, under Fe deprivation condition, PtrbHLH011 expression is inhibited, which is likely to activate Fe deficiency signaling and mechanisms to improve root-to-shoot Fe transport (Fig. 6). To further support this hypothesis, how these PtrbHLH011 targets affect Fe deficiency signaling and root-to-shoot Fe transport in poplar remains to be elucidated.

Fe deprivation has been reported to disrupt the photosynthetic machinery, resulting in the generation of light-dependent reactive oxygen species (ROS) and leaf photosensitivity (Akmakjian et al., 2021). Consistently, we observed significant GO enrichment of oxidation-reduction and superoxide metabolic processes in genes that were upregulated in Fe deprivation-treated poplar leaves (Fig. S3E). Since phenols and flavonoids are important antioxidant metabolites in plants, downregulation of PtrbHLH011 under Fe deprivation appears to protect plants from oxidative stress by reversing the repression of the expression of chalcone synthases (e.g. *CHS1* and *CHS5*) and increasing flavonoid biosynthesis (Fig. 6). Given the role of ferritins in oxidative stress response (Ravet et al., 2009), regulation on *FER1* expression is probably another contributor.

In summary, our results reveal a regulatory mechanism that controls SCW biosynthesis in response to environmental iron availability. Our mechanistic insights may suggest strategies of manipulating PtrbHLH011 expression and/or its regulatory relationships with downstream targets to enhance biomass production, nutrient usage and stress resistance of bioenergy crops under marginal conditions.

## Materials and Methods

### Plant materials and growth conditions

To generate PtrbHLH011 overexpression lines, full-length *Potri.005G113400* CDS (*P.trichocarpa* v4.1; Phytozome) was synthesized and cloned into the pTwist ENTR Kozak vector (Twist Bioscience). The gene was then subcloned into the Gateway binary destination vector pMDC32 using LR Clonase II recombination (Invitrogen) for overexpression. The resulting overexpression cassette is driven by the CaMV 35S promoter. The binary transformation vector was then transformed into *Agrobacterium* strain GV3101. The *P. tremula* x *P. alba* 717-1B4 genotype was transformed using a modified *Agrobacterium*-based method (Tsai et al., 1994). Shoots regenerated from isolated calli were tested by PCR (primers are listed in Table S1) to verify the presence of the transformed construct. Empty vector-transformed plants were used as controls. Plants were propagated in a greenhouse maintained at 20-24°C and a day length of 16 hours.

### Hydroponic treatments

Clonal cuttings of *Populus trichocarpa Nisqually-1* were grown in a hydroponic growth system in a greenhouse at 20-24°C and a day length of 16 hours. Woody cuttings of uniform size were rooted in water for 14 days and then in modified Hoagland’s solution (500 µM KNO_3_, 500 µM Ca(NO_3_)_2_, 100 µM NH_4_NO_3_, 200 µM MgSO_4_, 100 µM KH_2_PO_4_, 50 µM H_3_BO_3_, 10 µM MnSO_4_, 0.2 µM CuSO_4_, 0.5 µM Na_2_MoO_4_, 1 µM ZnSO_4_, 10 µM Fe, Na-EDTA, pH 5.6) for acclimation. The nutrient solution was renewed every 7 days throughout the experiment. After 21 days of acclimation in hydroponics, the roots were thoroughly washed with Milli-Q water and 20 mM EDTA solution, the plants were then randomized and transplanted into modified Hoagland’s solutions containing different Fe, Na-EDTA concentrations for treatment: Ctrl (10 µM Fe, Na-EDTA) and Fe deprivation (0 µM Fe, Na-EDTA). After 21 days of treatment, leaf and stem samples were collected for gene expression and cell wall analysis.

### Histological analysis

The internodes of the same age or treatment time were collected and cut into 80 µm thick sections using a cryostat (Leica CM1950). Sections were stained with 2% phloroglucinol-HCl or 0.02% toluidine blue O. The stained sections were imaged using the Odyssey M Imaging System (LI-COR) and the LMD7 Laser Microdissection microscope with a 10x objective (Leica). Cell wall thickness of over 40 xylem fiber cells was measured for each sample using LAS X software (Leica).

### Phylogenetic analysis

For reconstruction of the relatedness between poplar and Arabidopsis homologs, the PtrbHLH011 amino acid sequence was used to search the *Populus trichocarpa* v4.1 (Phytozome) and Arabidopsis thaliana TAIR10 with Blastp. The top matches were used to build a multiple sequence alignment using the CIPRES web portal (Miller et al., 2010) with MAFFT on XSEDE (v. 7.490) (Katoh and Standley, 2013). The IQ-TREE (Trifinopoulos et al., 2016) web server was used for construction of a phylogenetic tree under maximum likelihood using ultrafast bootstrap (UFBoot) (Hoang et al., 2018); the best-fit model according to BIC was JTT+F+I+G4 (Kalyaanamoorthy et al., 2017). For reconstruction of the relatedness of PtrbHLH011 homologs across land plants, FastTree (Price et al., 2010) with default parameters (JTT+CAT substitution model and 1000 bootstraps for a Shimodaira-Hasegawa test) was used. Trees were visualized using Interactive Tree Of Life (iTOL) and rooted at the midpoint.

### X-ray fluorescent and scattering microscopy

For XRF of leaf sections, freshly collected leaves were dissected into tissue pieces a few mm in size, rinsed with deionized water, and fixed in 4% paraformaldehyde (PFA) solution. The tissues were then sectioned to a thickness of 20 μm by using a cryostat (Leica CM1950). Sections were then placed on Silicon Nitride (SiN) membrane windows and air dried. All sections were examined for quality under the light microscope and those with best integrity were selected for scanning. Room temperature XRF measurements were performed on the 5-ID SRX beamline (Nazaretski et al., 2022) at BNL with a step size of 0.5 μm and a dwell time of 0.1 s. An AXO standard (AXO, Dresden GmbH) was used for XRF quantification. The spectra were analyzed and quantified using the open software PyXRF (Li et al., 2017) to create elemental distribution maps for each element.

Scattering-based tomographic imaging was used to examine the stem tissues at the LiX beamline at BNL (Yang et al., 2022). Data acquisition and processing were performed as previously described (Yang, 2024). The spatially resolved angular scattering intensity profile was divided into a constant background and an azimuthal-angle-dependent component. The ratio between the integrated intensity of these two components was calculated to represent the cellulose crystallinity index (CI). X-ray fluorescence data were acquired simultaneously with scattering data using a single-element silicon drift detector. The emission spectra were decomposed into contributions from known elements using non-negative least square fitting based on calculated characteristic emission peaks. The self-absorption by the sample was corrected during tomographic reconstruction using the algorithm developed by Ge et al (Ge et al., 2022).

### Anthocyanin, phenolics and total lignin measurements

Anthocyanin content of poplar leaf tissue was quantified as previously described (Nakata and Ohme-Takagi, 2014) with modifications. Briefly, 100 mg of leaf tissue from 4-month-old poplar plants was collected and ground using CryoMill (Retsch). The ground tissues were mixed with 1 ml of methanol with 1% HCl and extracted with rotation overnight at 20°C, followed by centrifugation to recover the supernatant. To quantify the anthocyanin content, 200 µl of supernatant of each sample was transferred to a 96-well plate and the absorbances at 530 nm and 637 nm were measured using the Spark Microplate Reader (Tecan). Each sample was analyzed with four technical replicates.

For total soluble phenolics measurement, ultraviolet-visible (UV-Vis) spectrophotometry and high-performance liquid chromatography-mass spectrometry (HPLC-MS) approaches were used. Soluble phenolics were extracted from approximately 100 mg of ground leaf tissue by incubating with 1 ml of 80% methanol overnight at 4°C. For UV-Vis spectra, the phenolic extract was diluted 30 times with methanol. The UV-Vis absorption spectra from 200 to 700 nm (1 nm resolution) of 150 µl of the diluted extractions were measured in UV-specific 96-well plate (Corning) using the Spark Microplate Reader (Tecan). 150 µl of methanol was measured as the background reference for reading normalization. For each sample three biological and three technical replicates were analyzed. HPLC-MS was performed as previously described (Dwivedi et al., 2024). The phenolic extracts were dried in a speed vacuum (Labconco) and digested in 2N HCl for 2 hours at 95°C. The acid digested samples were further extracted with 200 μl of water-saturated ethyl acetate. 100 μl of extracted samples were dried and redissolved in 200 μl of 80% methanol. 1 μl of the sample was injected into an UHPLC-MS system (ThermoFisher Scientific) and resolved with a reverse phase C18 column (Luna, 150 × 2.1 mm2, 1.6 μm, Phenomenex).

For total lignin measurement, Cell wall residues (CWRs) were prepared following the method described previously (Dwivedi et al., 2024). The total lignin content was measured using the acetyl bromide method (Foster et al., 2010). Approximately 10 mg of CWRs were incubated with 1 ml of 25% (v/v) acetyl bromide in glacial acetic acid at 50°C for 3 hours, with gentle mixing every 15 minutes. After incubation, the mixture was cooled on ice for 15 minutes and then diluted with 5 ml of glacial acetic acid. To neutralize the solution, 300 μl of this diluted solution was combined with 400 μl of 2 N NaOH and 300 μl of freshly prepared 0.5 M hydroxylamine hydrochloride. Next, 200 μl of the neutralized solution was transferred to UV-transparent 96-well plates (Corning, Kennebunk), and the absorbance at 280 nm was recorded. An extinction coefficient of 18.21 g−1 cm−1 was used to calculate the total lignin content.

### Protoplast transfection, subcellular localization and transient transactivation assays

Poplar mesophyll protoplasts were isolated and transfected as described previously (Xie et al., 2018b). Protoplasts were isolated from fully expanded leaves of two-month-old *Populus tremula* x alba 717-1B4 plants growing in magenta boxes.

For subcellular localization analysis, the cDNA of PtrbHLH011 was cloned into a transient expression vector (Xie et al., 2020) for the C-terminal YFP fusion. 5 µg of this plasmid was co-transfected with 5 µg of mCherry-VirD2NLS plasmid (Lee et al., 2008) into 100 µl of protoplast suspension (~100,000 cells). After 16 hours incubation under weak light at 23°C, protoplasts were collected and imaged with a Leica TCS SP5 confocal microscope equipped with 488 and 543 nm laser lines to excite YFP and mCherry, respectively. The emission bandwidth for YFP and mCherry was 500-530 nm and 580-620 nm, respectively. Images were processed using LAS X software (Leica).

Transient transactivation assays were performed as described(Xie et al., 2020). A total of 10 µg of effector, reporter, and/or transactivator plasmids were co-transfected into 100 µl of protoplast suspension (~100,000 cells). 100 ng of 35S:Luciferase plasmid was co-transfected for each reaction to normalize GUS activity. After 18 to 20 hours incubation in the dark at 23°C, protoplasts were collected and lysed to measure GUS and luciferase activities using the Synergy Neo2 multimode plate reader (BioTek). GUS activity in individual samples was normalized against luciferase activity (GUS/LUC). Three replicates were performed for statistical calculations.

### Transient ChIP-seq experiment and data analysis

Transient ChIP-seq was performed as previously described (Tadesse et al., 2024) using protoplasts isolated from *Populus tremula* x alba 717-1B4. Briefly, PtrbHLH011 was fused with the 10xMyc tag in a transient expression vector and transfected into 200 µl of poplar protoplast suspension (~200,000 cells). After 14 hours incubation at 23°C, protoplasts were collected for the µChIP experiment. Three biological replicates were prepared. ChIPed DNA and Input DNA were purified and concentrated using the MinElute PCR Purification Kit (Qiagen). Sequencing libraries were prepared using the ThruPLEX DNA-Seq Kit and DNA Unique Dual Index Kit (Takara) according to the manufacturer’s manual. 16 libraries were pooled for paired-end 100 bp sequencing using the DNBSEQ instrument (BGI). Approximately 20 – 30 M reads were obtained for each sample.

Clean reads were obtained by filtering raw reads to remove adapter sequences and low-quality reads. Due to the use of *Populus tremula* x *alba* 717-1B4 protoplasts, clean reads were mapped to the reference genome of *Populus tremula* and *Populus alba* (www.aspendb.org), respectively, using the BWA-MEM package (v0.7.17.2) to achieve the best mapping efficiency (Liu et al., 2014). Mapped data from ChIP samples and Input samples from three biological replicates with >80% correlation was then pooled for peak calling using MACS2 (v2.2.7.1) with *P* < 0.001. The ChIPseeker package (v1.18.0) was then used to annotate narrow peaks identified by MACS2. The gene lists from the *Populus tremula* and *Populus alba* mapping were combined by removing duplicate genes.

### RNA extraction, qPCR and RNA-seq analysis

Total RNAs were isolated from leaf and stem tissues using the RNeasy Plant Mini Kit (Qiagen) and treated with on-column DNase (Qiagen) to remove genomic DNA contamination. RNA quality was assessed using the 2100 Bioanalyzer (Agilent). For quantitative RT-PCR (qRT-PCR), cDNAs were synthesized using the oligo(dT) primer and RevertAid Reverse Transcriptase (Thermo Fisher Scientific) according to the manufacturer’s instructions. Primers for qRT-PCR are listed in Table S1.

For Fe deprivation RNA-seq, library construction and sequencing were performed by the Joint Genome Institute (JGI) to obtain paired-end reads of 150 bp. For *PtrbHLH011* OE RNA-seq, library construction and sequencing were performed by Beijing Genomics Institute (BGI) to obtain paired-end reads of 150 bp. Three biological replicates were initially used for library preparation and sequencing. But, because of the failures of library preparation, only two replicates were sequenced for Fe deprivation treated 3^rd^ internode and *PtrbHLH011* OE stem. After removing adapters and low-quality sequences, clean reads were aligned to the *Populus trichocarpa* genome v3.1 (Phytozome) using HISAT2 (v2.2.1). *Populus trichocarpa* genome v3.1 was used to be comparable with the genome information for the analysis of transient ChIP-seq data. Transcripts were then assembled using StringTie (v2.2.1). DEGs were identified using DESeq2 (v2.11.40.7) with the criteria of fold change ≥ 2 and *P* < 0.05. GO enrichment analysis was performed using the tool available on the PlantRegMap website (https://plantregmap.gao-lab.org/).

### DNA motif analysis

To predict the DNA motifs bound by PtrbHLH011, the sequences of the promoters tested in the transactivation assay were analyzed using the STREME (Sensitive, Thorough, Rapid, Enriched Motif Elicitation) function of the MEME suite (v5.5.2) (Bailey et al., 2015). Enriched DNA motifs in validated PtrbHLH011 targets (*PtrWND2B*, *PtrMYB92*, *CHS1*, *CHS5*, *YSL3*, *PYE*, *FER1* and *ZIP4*) were identified. Promoter sequences of genes that were not repressed by PtrbHLH011 (*ZIP6*, *TT3*, *PtrMYB3*, *SND2-1*, *SND2-2*, *MYB5* and *MYC2*; Fig. S7A) were used as control sequences. Promoter sequences are listed in Table S2.

### Protein purification and near-infrared fluorescence electrophoretic mobility shift assay (EMSA)

PtrbHLH011 was codon optimized, cloned into the pET-29b(+) vector for C-terminal 6xHis tag fusion, and transformed into *E. coli* BL21 (DE3) cells. Bacteria cells were grown in ZYM-5052 autoinduction medium (VWR) at 37°C to OD_600_ of 0.6 and then autoinduced at 20°C for overnight. Cells were collected and lysed using the B-PER Bacterial Protein Extraction Reagent (ThermoFisher Scientific) with lysozyme and benzonase. We observed that PtrbHLH011 is mainly in the insoluble pellet. To obtain soluble PtrbHLH011, the insoluble pellet was resuspended in the solubilization buffer (1% n-octyl β-d-glucopyranoside, 30 mM Tris pH 8.0, 10 mM sodium phosphate pH 8.0, 300 mM NaCl, 5% glycerol and 0.1mM DTT). PtrbHLH011-6xHis proteins were purified with Ni-NTA column and PtrbHLH011 proteins were then released by TEV protease cleavage.

EMSA was performed as previously described (Tadesse et al., 2024) with modifications. DNA oligonucleotides with 5’ IRDye 800 labeling were synthesized by Integrated DNA Technologies (IDT). Equal amounts of forward and reverse DNA oligonucleotides were annealed into a double-stranded DNA probe. 500 ng of purified protein and 2 μl of 5nM DNA probe were incubated for 20 min at room temperature. In parallel, the same amount of DNA probe was incubated with the protein buffer as the negative control. For competition assays, 10x, 50x or 100x unlabeled DNA probe was added. The reaction mixtures were then resolved in 6% DNA retardation gel (Novex) by electrophoresis at 100 V for 1 hour. The gel was then scanned using the Odyssey M Imaging System in the 800 nm channel (LI-COR).

## Data Availability

The RNA-seq and transient ChIP-seq data that support the findings of this study are openly available on Gene Expression Omnibus (GEO; https://www.ncbi.nlm.nih.gov/geo/), reference no: GSE265823 and GSE265825.

## Accession numbers

Sequence data can be found under the following PHYTOZOME accession numbers: (*P.trichocarpa* v4.1): PtrbHLH011 (Potri.005G113400), PtrWND2B (Potri.002G178700), PtrMYB92 (Potri.001G118800), PtCesA4 (Potri.002G257900), CHS1 (Potri.014G145100), CHS5 (Potri.003G176800), YSL3 (Potri.012G024700), PYE (Potri.015G142700), FER1 (Potri.016G124900), PtIRT3 (Potri.018G053300), ZIP6 (Potri.009G074100), TT3 (Potri.002G033600), PtrMYB3 (Potri.001G267300), SND2-1 (Potri.011G058400), SND2-2 (Potri.007G135300), MYB5 (Potri.001G005100) and MYC2 (Potri.001G142200) and eIF-5A (Potri.018G107300).

## Supplementary Data

Figure S1. Hydroponic iron (Fe) deprivation treatment of poplar plants.

Figure S2. Additional biological replicates of the histological analysis in Fig. 1A-D.

Figure S3. RNA-seq data analysis of poplar treated with iron (Fe) deprivation.

Figure S4. Phylogenetic reconstruction of PtrbHLH011 homologs from land plants shows that PtrbHLH011 is the close homolog of AtbHLH011.

Figure S5. Images of additional *PtrbHLH011* overexpression (*PtrbHLH011* OE) lines and phloroglucinol-HCl staining of the 3^rd^ internode cross-sections.

Figure S6. The reduction of total leaf soluble phenolic content in transgenic poplar with *PtrbHLH011* overexpression (*PtrbHLH011* OE).

Figure S7. RNA-seq and transient ChIP-seq data analysis to identify putative PtrbHLH011 targets.

Figure S8. Transient transactivation assay of putative targets not directly regulated by PtrbHLH011.

Table S1. Primers used in this study.

Table S2. Promoter sequences for motif discovery in Fig. 5B.

Data S1. Significantly downregulated and upregulated genes by comparing transcriptomes of Fe deprivation and control condition of the 3^rd^ internode.

Data S2. Significantly downregulated and upregulated genes by comparing transcriptomes of Fe deprivation and control condition of the leaf.

Data S3. Significantly downregulated and upregulated genes by comparing transcriptomes of *PtrbHLH011* OE leaf and empty vector leaf.

Data S4. Significantly down-regulated and up-regulated genes by comparing transcriptomes of *PtrbHLH011* OE stem and empty vector stem.

Data S5. Putative PtrbHLH011 targets identified by integrating RNA-seq and transient ChIP-seq data.

## Acknowledgments

We thank Katherine Fedotov at Stony Brook University for preparing leaf cross-sections for x-ray fluorescent microscopy experiment. This work was supported by the U.S. Department of Energy, Office of Science, Office of Biological and Environmental Research, as part of the Quantitative Plant Science Initiative (QPSI) at Brookhaven National Laboratory (BNL). The Fe deprivation RNA-seq (proposal: 10.46936/10.25585/60001386) was conducted by the U.S. Department of Energy Joint Genome Institute (https://ror.org/04xm1d337), a DOE Office of Science User Facility, is supported by the Office of Science of the U.S. Department of Energy operated under Contract No. DE-AC02-05CH11231. Flavonoid and lignin analysis was partially supported by the DOE, Office of Science, Office of Basic Energy Sciences, specifically the Physical Biosciences program of the Chemical Sciences, Geosciences and Biosciences Division under contract no. DE-SC0012704 (to C.-J.L.) and by the Joint BioEnergy Institute, one of the Bioenergy Research Centers of the US DOE, Office of Science, Office of Biological and Environmental Research, through contract DE-AC02-05CH11231 between Lawrence Berkeley National Laboratory and the U.S. Department of Energy. Work at the Molecular Foundry was supported by the Office of Science, Office of Basic Energy Sciences, of the U.S. Department of Energy under Contract No. DE-AC02-05CH11231 (CEB-H). The X-ray measurements were conducted at NSLS-II of Brookhaven National Laboratory, a National User Facility supported in part by the U.S. Department of Energy under contract number DE-SC0012704. The LiX beamline is part of the Center for BioMolecular Structure (CBMS), which is primarily supported by the National Institutes of Health, National Institute of General Medical Sciences (NIGMS) through a P30 Grant (P30GM133893), and by the DOE Office of Biological and Environmental Research (KP1605010). This research used the confocal microscope of the Center for Functional Nanomaterials (CFN), which is a U.S. Department of Energy Office of Science User Facility, at Brookhaven National Laboratory under Contract No. DE-SC0012704.

## Author Contributions

M.X. conceived and designed the research; M.X. wrote the manuscript; J.C. and K.B. edited the manuscript; D.T., Y.D., N.D., D.K. and K.S. performed experiments; L.Y., Y.Y. and K.F. performed and analyzed X-ray experiments; C.E.B. performed phylogenetic analysis; M.X. and A.L. analyzed the data. All authors reviewed and approved the final version of the manuscript for publication.

